# Disentangling the role of chromosome inversions in the structure of flat oyster (*Ostrea edulis*): origin, adaptation and phylogeography

**DOI:** 10.1101/2025.06.12.659260

**Authors:** I. M. Sambade, V. Hlordzi, A. Blanco, M. Hermida, M. Vera, S. Culloty, T. Bean, S. A. Lynch, P. Martínez

**Affiliations:** University of Santiago de Compostela; University College Cork; Roslin Institute

**Author notes:** Corresponding author: Paulino Martínez.

## Abstract

Chromosomal inversions are large-scale structural variants that disrupt recombination and contribute to population differentiation and local adaptation. In this study, we characterized the genomic features, frequency, and evolutionary significance of chromosomal inversions in the European flat oyster (*Ostrea edulis*), a species of ecological and commercial relevance. Using a low-density SNP array (∼5K), we obtained confident genotyping data across the native European distribution of the species. The breakpoints of the inversions at chromosome (C) 2, C5 and C8, spanning 27, 7 and 35 Mb, respectively, were precisely mapped by combining linkage disequilibrium in a large sample and long-read sequencing. This enabled us to identify the SNPs and genes within the inversions for comparative population genomics and functional evaluation. A north to south cline was observed for the three inversions, although with low-moderate correlation between them. Strong signals for divergent selection were detected for the three inversions, although with specific profiles suggestive of adaptive peaks around highly divergent SNPs across the inversion length. The three inversions seem to be old considering genetic diversity and differentiation between arrangements. Phylogeographic events and adaptation to the current environment seem underlying the structural pattern observed. Gene content and enrichment analyses within inversions suggest that these polymorphisms may play a role in the immune system and response to different stressors including parasite resistance. Our results provide new insights into the genomic architecture of adaptation in *O. edulis* and highlight the potential of structural variants as targets for conservation and selective breeding.

## Introduction

Chromosomal inversions, first discovered in *Drosophila* over a century ago (Sturtevant, 1920), represent a type of structural variant in which a DNA segment changes its orientation within the chromosome. These inversions have long been implicated in adaptation by ensuring the co-transmission of favourable allele sets as haplotypes through recombination suppression (supergenes; Dobzhansky, 1970; Reeve et al., 2023). Alleles in regions with reduced recombination are inherited as large units, amplifying the signals of forces that generate islands over extensive genomic regions (Berdan et al., 2023). This facilitates adaptation to specific environments that could reinforce reproductive isolation between diverging populations by accumulation of genetic incompatibilities, which can reduce hybrid viability or fertility and, consequently, drive speciation (Lowry & Willis, 2010; Zhang et al., 2022). The persistence of these polymorphisms can be explained by divergent selection, balancing selection, or a combination of different processes within the evolutionary history of species (Kirkpatrick & Barton, 2005; Faria et al., 2019). From a population genetics perspective, chromosomal inversions can play a crucial role in local adaptation to distinct habitats or environmental conditions (Noor et al., 2001) and have been frequently associated with environmental clines (Wellenreuther & Bernatchez, 2018). Understanding the maintenance and distribution pattern of these polymorphisms should be interpreted at the light of the evolutionary history of the species (Monteiro et al., 2024).

Inversions usually involve long chromosome tracts spanning up to 100LMb and collectively can account up to more than 50% of the genome (Wellenreuther & Bernatchez, 2018; Walkowiak et al., 2020). They can include the centromere (pericentric) or not (paracentric), but the latter have been more frequently reported. Additionally, chromosomal inversions can persist for thousands or even millions of generations, as for example the 900 kb inversion in humans originated approximately three million years ago (Stefansson et al., 2005). For decades, chromosomal inversions have been used to construct phylogenies and to study geographic and temporal cycles (Krimbas & Powell, 1992). Although their study declined with the rise of molecular genetics in the 1970s (Kirkpatrick, 2010), it has regained prominence with new genomic technologies and bioinformatic tools that provide precise information of the boundaries and genetic constitution structural variants and their frequency across the distribution range of species (Wellenreuther and Bernatchez, 2018; De Coster & Van Broekhoven, 2019). Recent studies have confirmed that inversions reduce recombination in their surrounding regions and can generate population structure, accumulating more genetic variation than non-inverted regions (Wellenreuther & Bernatchez, 2018).

Chromosomal inversions are common across a wide range of aquatic organisms and have been associated with adaptive variation in environments without obvious barriers to gene flow (Berg et al., 2016; Catanach et al., 2019; Pearse et al., 2019; Hollenbeck et al., 2022; Thorstensen et al., 2022). In molluscs, inversion polymorphisms have been associated with different ecotypes across the distribution range in *Littorina saxatilis* (Reeve et al., 2024), to a surface temperature water cline in *Pecten maximus* (Hollenbeck et al., 2022) or to an arginine kinase (*ark*) polymorphism putatively involved in shell size in *Littorina fabalis* (Le Moan et al., 2022). Bivalves represent a model for investigating the relationship between genome architecture and adaptation, as there is substantial evidence of local adaptation across heterogeneous environments (Hollenbeck et al., 2022), along with a growing body of data documenting an exceptional degree of structural variation in their genomes (Calcino et al., 2021).

The European flat oyster, *Ostrea edulis* (Linnaeus, 1758), is a bivalve mollusc of great ecological and economic relevance. With a history of human exploitation dating back to the Mesolithic era, this species has been a staple in the seafood market for centuries and plays a vital role in marine ecosystems by contributing to water filtration and providing habitat for marine life (Astrup et al., 2019; Pogoda et al., 2022). Beyond its ecological functions, *O*. *edulis* is also a key species in aquaculture, cultivated for both commercial purposes and biodiversity restoration initiatives (Albentosa et al., 2023). Oyster aquaculture not only helps meet global seafood demand but also reduces pressure on wild stocks by providing a sustainable production method. It supports biodiversity through reseeding efforts and allows for controlled growth and to control pathogens through breeding programs (Zu Ermgassen et al., 2023).

*Ostrea edulis* currently distributes from Northern Europe, where it is found in the North Sea and the Baltic Sea, to Southern Europe, along the Atlantic coast of France and Spain (FAO, 2022; Pouvreaeu et al., 2023). In the Mediterranean Sea, it occurs in specific areas such as the Adriatic, Aegean and Black seas (Acarli et al., 2011; Stagličić et al., 2020), as well as the Mar Menor lagoon in Spain, where a population has been reintroduced to aid controlling pollution (Albentosa et al., 2023). This species shows a complex structure across its natural distribution due to phylogeographic events, historical translocations, restricted gene flow due to oceanic dynamics (fronts, currents), and local adaptation (Vera et al., 2016; Lapègue et al., 2022; zu Ermgassen et al, 2023; Monteiro et al., 2024). Early analyses with allozymes (Saavedra et al., 1993) indicated low geographic differentiation, although specific loci, such as arginine kinase (*ark*), exhibited a pronounced cline across the sampled Atlantic area. Microsatellite data supported three main regions across the Northeast Atlantic mainly associated with the Ushant front (Southern region) and in the north two additional regions associated with the English Channel, the British Isles and the North Sea (Vera et al., 2016). Four Atlantic mtDNA haplogroups have been identified using ancient DNA, one showing a patchy distribution from the North Sea to the Atlantic coast of France (Hayer et al., 2021). The other three haplogroups would be restricted to narrow geographic ranges, suggesting adaptation to local environmental conditions or oceanic barriers to gene flow, but also related to the glaciations along Pleistocene involving glacial refugia and post-glacial expansion (Hayer et al., 2021). Genetic analyses with nuclear markers supported an isolation by distance model but surprisingly grouping North Sea populations with those from the Eastern Mediterranean and Black Sea (Diaz-Almela et al., 2004; Lapègue et al., 2022).

High structural variation has been reported in flat oyster, including long inversions in three chromosomes (Sambade et al., 2022; Lapègue et al., 2022; Monteiro et al., 2024) of its karyotype (n = 10, Thiriot-Quiévreux, 1984). Specifically, inversion of C8 accumulates the most relevant outliers associated with resistance to the parasite *Bonamia ostreae* (Vera et al., 2019; Sambade et al., 2022). Moreover, these inversions seem to follow a geographical pattern, which could result from phylogeographic events or be related to adaptation to specific biotic or abiotic environmental factors (Monteiro et al., 2024).

The present study aimed at characterizing the three large chromosome inversions of *O*. *edulis* at genomic level to understand their role on population structure and adaptation of the species across its distribution range. We precisely established the limits of the inversions using long-read sequencing combined with a linkage disequilibrium approach and evaluated the distribution of arrangements within and among populations across the distribution range of the species to ascertain their role in adaptation at the light of phylogeographic events. This information will be useful both for management of natural resources as well as for implementing sustainable production of flat oyster.

## Materials and Methods

### Sampling

Population genomics analyses were conducted on 343 wild oysters collected in 13 shellfish beds (Table 1; Fig 1). Some samples came from previous studies or projects in flat oyster, while others were collected for this study to cover, as far as possible, the distribution range of the species. This sampling helped to refine population structure of the species at the light of the main chromosome inversions identified in the species (Sambade et al., 2022; Lapègue et al., 2022; Monteiro et al., 2024; Table 1). Moreover, 384 oysters collected from May to September 2023 in two close Irish populations (∼125 nautical miles; Lough Swilly, LW and Galway, GAL; Fig. 1) showing very low genetic differentiation between them (see Results), were employed to explore putative deviations from Hardy-Weinberg (HW) equilibrium due to selection and for delimiting the boundaries of the inversions through linkage disequilibrium (LD).

**Figure 1.**
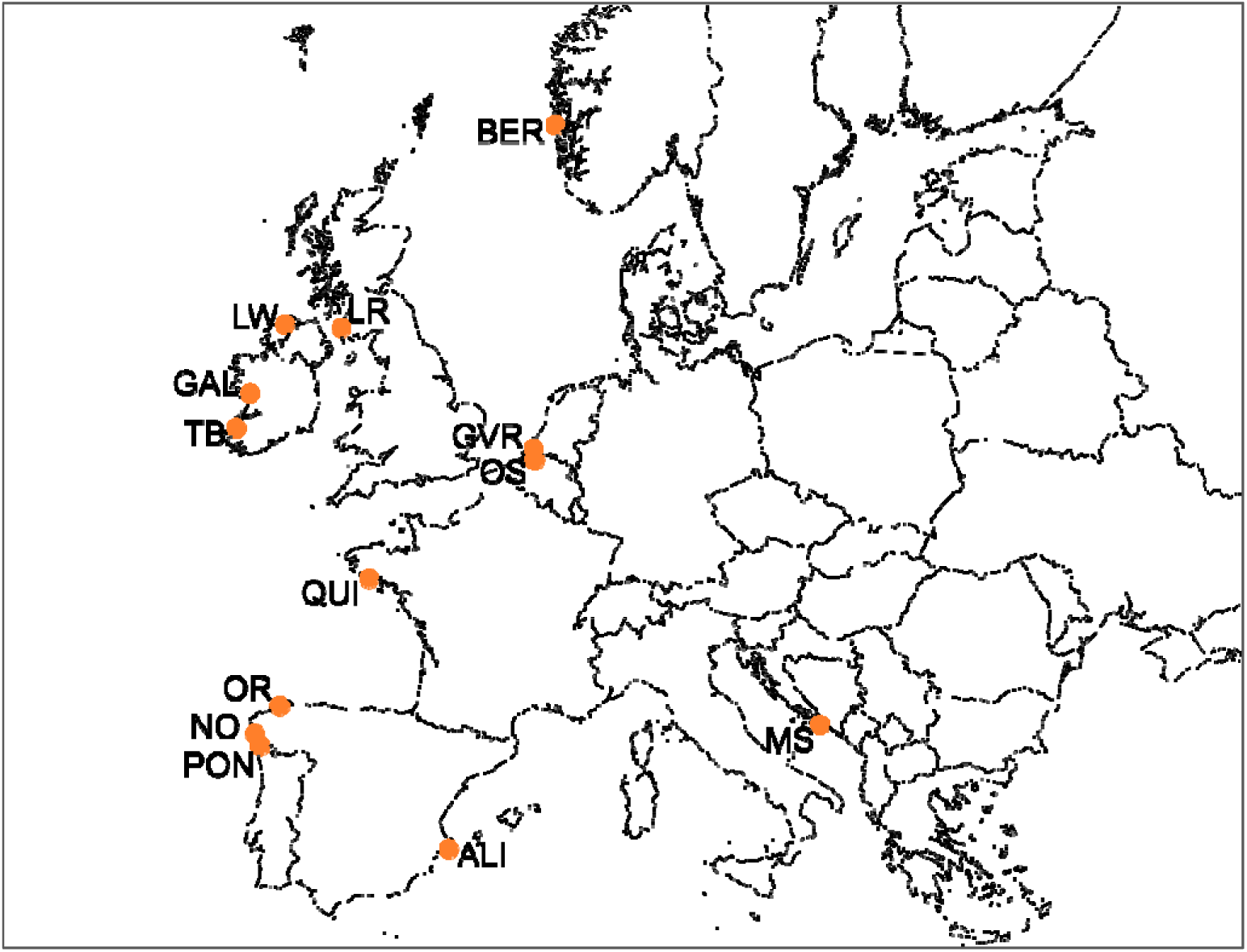
Sampling locations of *O*. *edulis* in this study. Sampling codes from Table 1.

**Table 1.** Main features of the *O*. *edulis* shellfish beds.

| Location | Code | Country | N | Sample collection |
| --- | --- | --- | --- | --- |
| Bergen | BER | Norway | 30 | 1 |
| Grevelingen | GVR | The Netherlands | 21 | 1 |
| Oosterschelde | OS | The Netherlands | 28 | 1 |
| Loch Ryan | LR | Scotland | 16 | 2 |
| Lough Swilly | LW | Ireland | 33 (192) | this study |
| Tralee Bay | TB | Ireland | 16 | 2 |
| Galway | GAL | Ireland | 37 (192) | this study |
| Quiberon | QUI | France | 15 | 2 |
| Ortigueira | OR | Spain | 16 | 2 |
| Noia | NO | Spain | 33 | 3 |
| Pontevedra | PON | Spain | 33 | 3 |
| Alicante | ALI | Spain | 30 | this study |
| Mali Ston Bay | MS | Croatia | 35 | 3 |

1. Sambade et al. (2022), 2. Vera et al. (2019), 3. Vera et al. (2016), 4. Šegvić-Bubić et al. (2020); in parentheses the samples used to test HW and LD for the three inversions in LW and GAL.

### DNA extraction

Genomic DNA for population genomics was extracted from gill tissue using the E.Z.N.A.® Mollusc DNA kit (OMEGA Bio-Tek) following the manufacturer’s recommendations. Briefly, between 30 and 50 mg of fresh gill or stored in 100% ethanol were treated with a solution containing 20 μl of proteinase K and 700 μl of lysis buffer (CSPL) overnight, while gently shaking to mix thoroughly at 65°C. Samples were subjected to RNAse A treatment and to several buffer washes by centrifugation. DNA quantity and purity was measured with a Nanodrop spectrophotometer, and DNA concentrations were normalized for single nucleotide polymorphism (SNP) genotyping.

### SNP genotyping

Genotyping of flat oyster samples from Vera et al. (2019) and Sambade et al. (2022) was performed with a multispecies Affymetrix Axiom Custom SNP Array (∼15K SNPs for *O*. *edulis*; Gutiérrez et al., 2017). The remaining samples were genotyped with a subset of 4,741 SNPs of that array selected from population genomics data (Vera et al., 2019; Sambade et al., 2022). Selected SNPs were included in a multispecies Affymetrix Axiom SNP Array developed by the University of Santiago de Compostela in collaboration with Thermo-Fisher Scientific and Xenética Fontao SA according to the following criteria: i) Minimum Allele Frequency (MAF) > 0.05; ii) missing data < 0.2; and iii) homogeneous distribution across the *O*. *edulis* genome (Gundappa et al., 2022) (Table S1A). Average SNP genomic density was 4.9 SNPs / Mb, ranging between 2.8 SNPs / Mb in C9 and 6.9 SNPs/Mb of C5 and C6 (Fig. S1). Genotyping information for this set of 4,741 SNPs was also retrieved from the original 15K SNP array for the samples analysed by Vera et al. (2019) and Sambade et al. (2022), so all samples in this study were analysed with the same SNP dataset.

### Genomic characterization of chromosome inversions

To establish the boundaries of the main inversions in the flat oyster genome (Gundappa et al., 2022) detected in previous studies at chromosome C2, C5 and C8 (Sambade et al., 2022; Lapègue et al., 2022: LG1, LG6 and LG4, respectively; Monteiro et al., 2024: scaffold5, scaffold4, scaffold8; see Results), we followed a two-step approach.

#### Linkage disequilibrium

Firstly, LD was evaluated in a large sample of 384 oysters to obtain a rough estimation of the inversion breakpoints, considering the average SNP density (4.9 SNPs / Mb). Samples were collected from May to September 2023 in LW and GAL (Fig. 1, Table 1), a geographical region where previous studies suggested high polymorphism for the inversions (Vera et al., 2019; Lapègue et al., 2022, Monteiro et al., 2024).

*r*^2^ was first estimated using PopLDdecay (Zhang et al., 2019), and the resulting data was represented in scatterplots generated with ggplot2 package in R (Wichham, 2016), where *r*^2^ values were plotted against the physical distance between SNPs to evaluate the decline of LD across chromosomes. For chromosomes showing an unusual LD decay pattern, pairwise LD was calculated for all SNPs using PLINK 1.9b5 (Chang et al., 2015) without a maximum distance threshold. LD values were then visualized as heatmaps using the LDheatmap package in R (Shin et al., 2006), enabling us to establish the boundaries of inversions in a range of tens kilobases at C2, C5 and C8 (see Results).

Furthermore, LD variation across C2, C5 and C8 inversion lengths was analysed by inspecting LD profiles using a sliding window of ten consecutive SNPs, where *r*^2^ was averaged across all pairwise comparisons using PLINK 1.9b5, and the window shifted by one SNP per step. This information was also used to explore the correspondence between LD and genetic differentiation across the inversion lengths.

Since the estimated length of the inversions in our study did not match those reported by Lapègue et al. (2022) and Monteiro et al. (2024), two comparisons were performed: i) a BLAST (Altschul et al., 1990) was conducted to align the microsatellites used by Lapègue et al. (2022) against the reference genome from Gundappa et al. (2022); ii) Minimap2 (Li, 2018) was used to align the genome from Boutet et al. (2022) against that from Gundappa et al. (2022). The resulting PAF (Pairwise mApping Format) files were visualized using the R packages ggplot2 (Wickham, 2016) and pafr (Winter, 2020).

#### Long-read high-fidelity (Hi-Fi) PacBio sequencing

To precisely establish the breakpoints of inversions, PacBio HiFi whole genomic sequencing (WGS) was performed on six individuals with suitable genotypes for the inversions studied in the large Irish populations (LW and GAL) (see Results). Sequencing was performed at the Novogene platform using a 20X coverage according to the estimate flat oyster genome (935 Mb; Gundapa et al., 2022). Raw reads were filtered using the chopper software (De Coster and Rademakers, 2023) with the following parameters: i) minimum quality threshold 20; ii) minimum read length 5000 bp; iii) initial and final trimming 10 bp from each end. Initial metrics were evaluated for each sample, including the total number of reads, minimum and maximum lengths, average read length, and total read size to assess the quality of the processed data.

To establish the limits of C2, C5 and C8 inversions, we retrieved 1000 bp from both ends of each long read, which were then aligned to the *O. edulis* genome (Gundappa et al., 2022) using Bowtie2 (Langmead & Salzberg, 2012; Langmead et al., 2018). Those reads overlapping the breakpoints of the inversion should be broken in any of the two arrangements for each inversion, depending on the arrangement in the reference genome, and shifted far away at the other end of the inversion breakpoint. Reads, where initial and final 1000 bp ends mapped at least 1 Mb apart, were retained for further inspection. The fragments were aligned to the windows delimited through LD analysis using a custom script, to be further inspected using the Integrative Genomics Viewer (IGV) software (Robinson et al., 2011). Those fragments ending at the same point were inspected as the putative breakpoints of each inversion.

#### Validation of inversion breakpoints through PCR: a tool for genotyping inversions

To confirm the breakpoints identified, PCR primers at both ends for each of the three inversions were designed using the flanking sequences in the reference genome. Primers were designed using Primer3 (Untergasser et al., 2012) and special care was taken to avoid annealing out of the target regions considering the abundance of repetitive elements (see Results). PCR tools for the three inversions were validated using homokaryon and heterokaryon individuals using the information obtained from population genomics data (see Results). PCR was carried out in a thermal cycler (Applied Biosystems 2720) with the following program: initial denaturation at 95°C for 10 minutes; 35 cycles of 94°C for 45 seconds for denaturation, 58°C for C2 and C5 and 60°C for C8 for 50 seconds for annealing, and 72°C for 50 seconds for extension; followed by a final extension at 72°C for 10 minutes.

#### Origin of inversions

Once established the limits of the inversions, we looked for repetitive elements around the breakpoints suggestive of specific mechanisms involved in their origin. Patterns of variation at flanking regions can be informative on the origin and history of inversions (Orengo et al., 2019). The presence of repetitive elements (RE) was explored using RepeatMasker 2.0, while the presence of regions enriched in tandem repeats was explored using a custom script that split the regions in 100bp windows, which were then aligned with Bowtie2 to detect duplications. Repetitive elements have been suggested to underline the specific mechanism on the origin of inversions through ectopic recombination or breakage / joining mechanisms (Delprat et al., 2009; Villoutreix et al., 2021: Pascarella et al., 2022; Huang et al., 2025).

### Genetic structure at the light of inversions

Once delimited the limits of inversions, genetic diversity and structure was investigated across the distribution range of the species considering the structural variation (arrangements) and using different SNP datasets: i) all the 4,741 SNPs; ii) all SNPs excluding those mapping within the inversions (4,186 SNPs); iii) the subset of SNPs within each inversion: C2 (209), C5 (65), and C8 (151) (see Results).

Genetic diversity (He) and conformance to HWE and the sense and magnitude of the deviation from random mating were evaluated with the R package Genepop 4.7.5. Pairwise population differentiation (F_ST_; Weir and Cockerham, 1984) was estimated using R package Genepop 4.7.5 (Rousset, 2008) with the ‘Fst’ function. StructureSelector software (Li and Liu, 2018) was used to obtain K estimators and CLUMPAK outputs (Kopelman et al., 2015). This program determines the most likely number of genetic clusters (K), defined as the number of population genetic units, by analysing linkage disequilibrium and HWE under a Bayesian clustering approach. To achieve this, STRUCTURE (Pritchard et al., 2000) was run with K values ranging from X to Y, using Z iterations per run and a burn-in period of W iterations. The resulting STRUCTURE outputs were processed in StructureSelector to determine the most likely K based on ΔK (Evanno et al., 2005). Discriminant analysis of principal components (DAPC), a multivariant method to infer the number of clusters within a group of genetically related individuals, was employed as a complementary approach to disclose genetic structure in the studied samples. The Adegenet package function ‘dapc’ in RStudio was used (Jombart and Ahmed, 2011). A principal component analysis (PCA) from the matrix of genotypes was performed and then, a selected number of principal components (PCs) used as input for linear discriminant analysis (LDA). To determine the optimal number of PCs for LDA, a cross-validation process was implemented, and the PCs associated with the lowest Root Mean Square Error (RMSE) retained.

### Signals of selection

Deviations from Hardy-Weinberg equilibrium (HWE) for structural variants was inspected in the big Irish population (GAL+LW) to ascertain putative mechanisms underlying the maintenance of these polymorphisms in flat oyster (i.e. overdominance, divergent selection).

Also, genetic diversity and differentiation profiles were explored across the full length of each inversion, either using single SNPs or their average values across sliding windows of 5 / 10 SNPs with one SNP shift per step. Genes around the putative adaptive peaks identified within each inversion and the breakpoints were inspected to look for candidates suggesting a functional explanation on their origin and evolution.

Outliers deviated from the neutral genomic background were investigated with BAYESCAN v2.01 (Foll & Gaggiotti, 2008), which employs an F_ST_-based Bayesian method based on single SNP evaluations, a suitable method for the scenario explored (see Results). Bayescan was run using default parameters (i.e., 20 pilot runs; prior odds value of 10; burn-in of 50,000), 100,000 iterations and a sample size of 5,000.

The list of coding genes within each inversion was retrieved from the reference genome (Gundappa et al., 2021) and functionally annotated with Pannzer (Törönen and Holm, 2022). Functional enrichment of genes located within each inversion was obtained with R package GOfuncR version 1.14.0 (Grote, 2021).

## Results

### Genomic characterization of chromosome inversions

The two large Irish samples (GAL and LW) showed very low genetic differentiation (F_ST_ = 0.0074; P = 0.093), so they were pooled (384 individuals) to increase the statistical power for LD analysis. Using the 4,741 SNP panel, atypical patterns of LD decay with physical distance were detected in three chromosomes. Most chromosomes showed the usual L-shaped decay curve, where *r*^2^ values dropped below 0.2 at ∼100 kb (Fig. S2A), except C2, C5 and C8, where *r*^2^ average values remained high and significant across ∼25 Mb, ∼7 Mb, and ∼40 Mb physical distance, respectively (Figs. S2B, S2C and S2D).

Pairwise LD at C2, C5 and C8 was high and significant across regions delimited by specific SNPs (Fig. S3), suggesting large inversions disrupting recombination. The abrupt LD drop between SNPs at both ends enabled delimitation of genomic windows where the breakpoints might be located. For C2, LD extended from the start of the chromosome (between 1 bp and 779,719 bp) to ∼27.5 Mb (between 27,544,809 bp and 27,592,275 bp) (Fig. S3A); for C5, from ∼86.4 Mb (between 86,428,703 bp and 86,508,092) to 93.1 Mb (between 93,012,400 bp and 93,108,624 bp) (Fig. S3B); and for C8, from ∼33.9 Mb (between 33,908,317 bp and 35,041,583 bp) to 69.6 Mb (68,632,940 bp and 69,558,118 bp) (Fig. S3C).

LD variation across C2, C5 and C8 was also inspected by averaging *r*^2^ over 5 / 10 SNPs using sliding windows in the big Irish sample (Figs. S4A, B, C) and in all samples throughout the distribution range (Figs. S4D, E, F). Despite an irregular and similar saw-peak pattern was observed in both datasets, the lower LD values within each inversion were always high above the chromosome LD background. Accordingly, when heatmaps were generated separately for homokaryon individuals for each arrangement no LD was detected at C2, C5 and C8 (Fig. S5). These observations strongly support the presence of single long inversions at these three chromosomes (Fig. 2). Considering that the flat oyster karyotype is mostly constituted by biarmed chromosomes, especially the biggest ones (Thiriot-Quiévreux, 1984), it seems very likely that the C2 and C5 inversions are paracentric. The C8 could correspond to a submetacentric chromosome, assuming a rough correspondence between scaffold and chromosome size, but as for the other two chromosomes, a LD peak likely corresponding to the centromere, could also suggest a paracentric inversion for C8 (Fig. S4).

**Figure 2.**
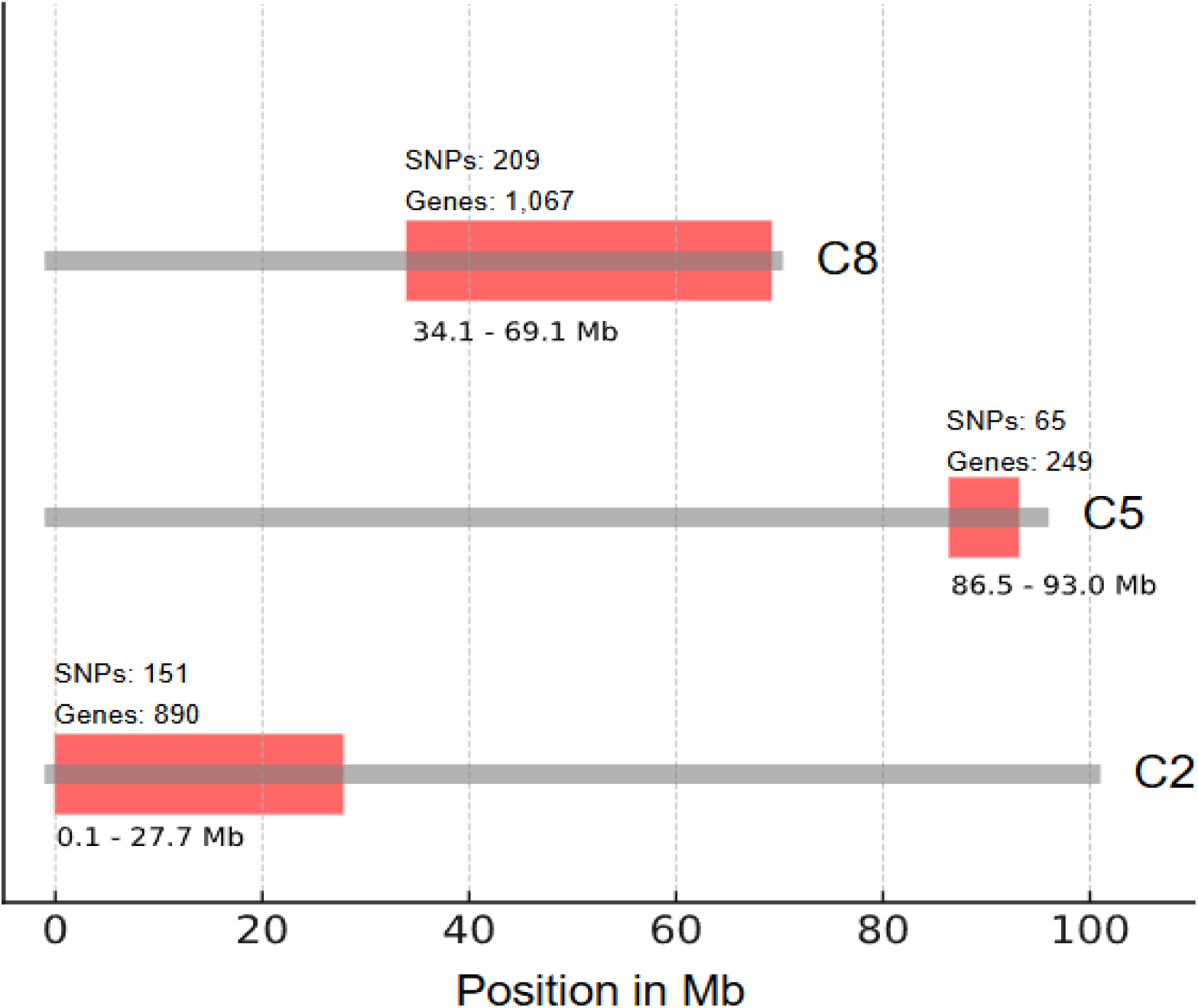
Main genomic features of the three chromosomic inversions in *O*. *edulis*.

The three inversions here detected corresponded to the linkage groups (LG)1, LG4 and LG6 reported by Lapègue et al. (2022) according to the blast performed with sequences of flanking microsatellites from the genetic map by Harrang et al. (2016) against the *O*. *edulis* reference genome (Gundappa et al., 2022). Minimap2 alignments between the Gundappa et al. (2022) and the Boutet et al. (2022) genomes showed a correspondence between scaffold4, scaffold5 and scaffold8 reported by Monteiro et al. (2024) with C5, C2 and C8. Despite the much lower number of markers in the study by Lapègue et al. (2022), the relative size of the inversions roughly matched to our findings; however, the inversions by Monteiro et al. (2024) showed remarkable differences at scaffold5 vs C2 (11.2 Mb vs 27.5 Mb) and scaffold8 vs C8 (24.5 vs 34.8 Mb). A ∼10Mb fragment of C8 was missing in the Monteiro et al. (2024) genome with respect to Gundappa et al. (2022), and the alignment of C2 vs scaffold5 was very fragmented including several opposite orientations (Fig. S6).

The six individuals from the big Irish sample selected for Hi-Fi sequencing included both arrangements for the three inversions, either in heterokaryon or homokaryon state according to population genomics information. SNPs within each inversion from the LD analysis (Table S1B) analysed with STRUCTURE program enabled an straightforward classification homokaryons or heterokaryons, showing a full or intermediate constitution for each cluster within each inversion (Fig. S7; Table S2). Total reads ranged from 876,960 (GAL_7) to 1,393,841 (LW_33) with average read lengths ranging from 6,675 (LW_33) to 11,635 (GAL_7), excluding GAL_173 that showed very poor sequencing data and was discarded for further analyses (Table S3). After quality filtering, reads ranged from 57.7% (LW_33) to 87.4% (LW_15). From this dataset, broken long reads with fragments separated by > 1 Mb were aligned to the LD windows in the reference and fragments perfectly aligning at both ends of each inversion were inspected. According to this information, C2 inversion spanned from 80,922 up to 27,676,307, C5 from 86,546,053 up to 93,001,597, and C8 from 34,069,607 bp up to 69,085,710 bp.

To confirm the location of the breakpoints and tuning a simple molecular tool for genotyping, PCR primers flanking breakpoints at both ends of each inversion were designed, and the banding patterns expected in agarose gels were compared with the population genomics information in a sample of ∼100 individuals (Fig. 3). Furthermore, the amplicon of each PCR was sequenced on an ABI3750 automatic sequencer (Applied Biosystems) and aligned to the genomic regions flanking the inversion breakpoints for confirmation. PCR results were consistent with STRUCTURE and sequencing information at both ends of C8 and at the 5’ end of C2 and 3’ end of C5, but the PCRs at the 3’ end of C2 and 5’ end of C5 inversions failed to amplify. For the 3’ end of C2 and 5’ end of C5, the interval established through LD was used for further analyses (Figs. 3A and 3B).

**Table 2.**
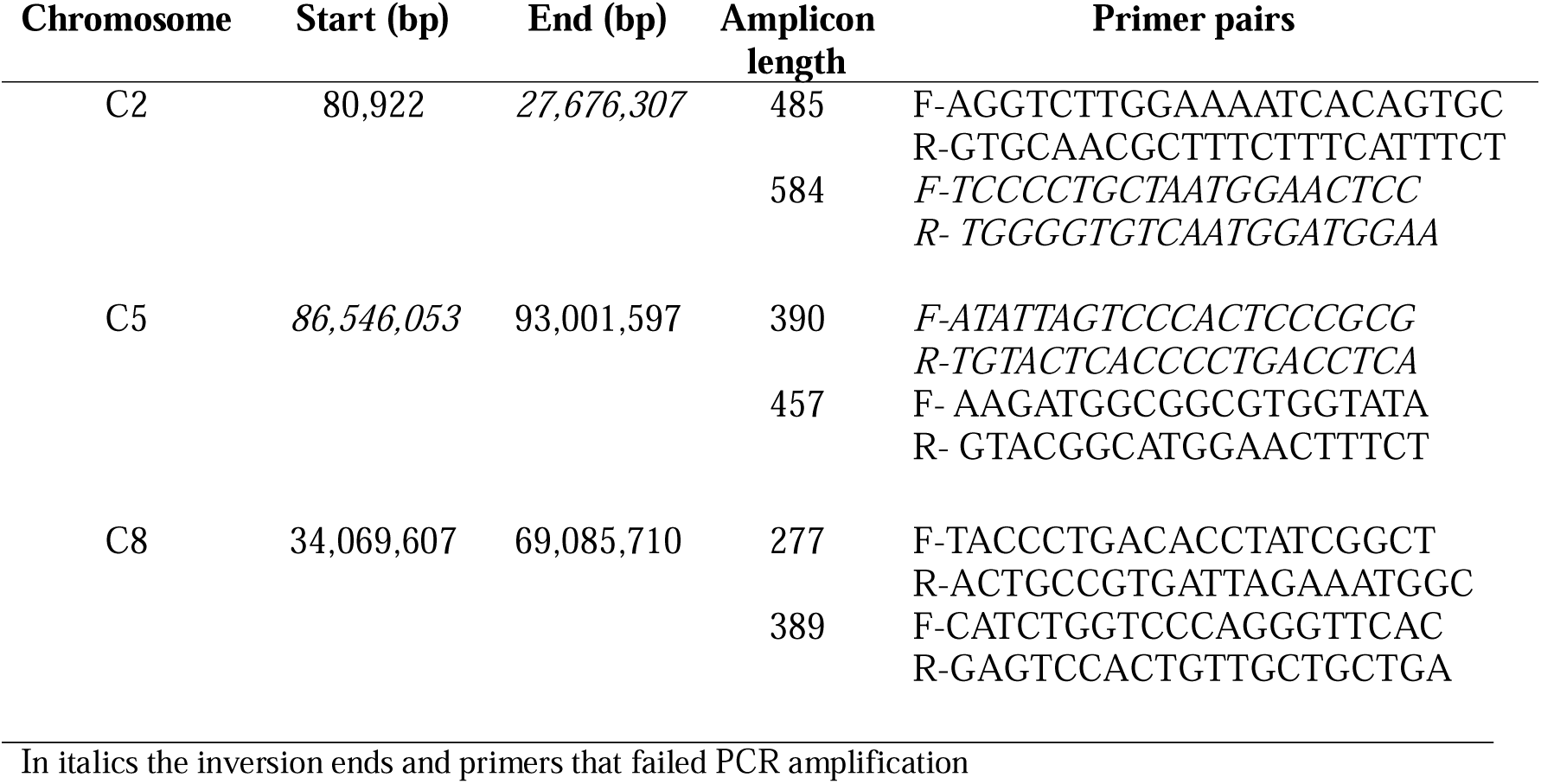
Genomic features of the inversions at C2, C5 and C8 of *O. edulis* and PCR settings for genotyping.

**Figure 3.**
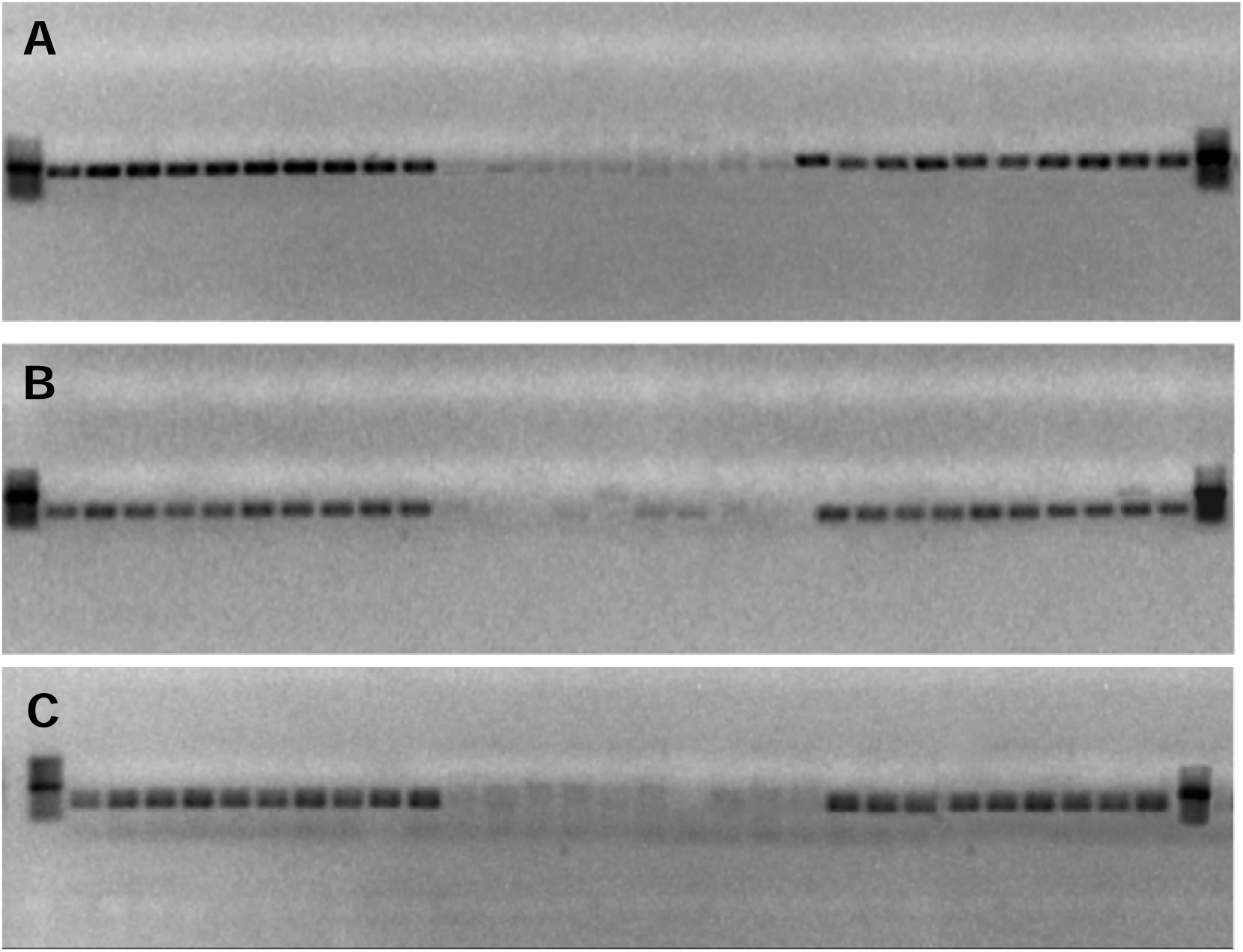
Agarose gels of representative *O*. *edulis* heterokaryon and homokaryon individuals for both arrangements at: A) C2 5’end, B) C5 3’ end, and C) C8 5’ end using a PCR tool; 100bp ladder at both ends; lanes 1-10: homokaryon for one arrangement; lanes 11-20: homokaryons for the opposite arrangement; lanes 21-30: heterokaryons.

The breakpoints at both ends of each inversion were inspected to look for repetitive elements (REs) that could be involved on their origin by ectopic recombination or break / joining events. In the four confirmed breakpoints (5’ C2, 3’ C5 and 5’ and 3’ C8), a LINE element flanked the breakpoints at one side, and at the other a DNA/hAT (5’ C2 and 3’ C8) or a PLE/Naiad (3’C5 and 5’ C8) were found (Table S4); furthermore, the distance between the end of one element and the start of the other was short (5’ C2: 9.4 kb; 3’ C5: 1.4 kb, overlapping 173 bp with PLE/Naiad; 5’ C8: 1.3 kb, 3’ C8: 2.5 kb). On the other hand, using 100 bp windows around the flanking regions, we could detect duplicated segments not related to transposable elements. Extensive duplications were observed in C2 at 71,500 – 74,400 bp and at 27,788,100 – 27,792,500 bp; and in C5 at 86,497,300 – 86,503,200 bp and 93,004,200 – 93,006,100 bp.

Gene Ontology (GO) terms of annotated genes within inversions were investigated to look for gene clusters that could provide functional insights into the adaptive role of inversions (Table S5). Among the 890 annotated genes at C2 inversion, biological process was enriched on neural detection and transmission of external and abiotic stimuli, including epidermal cell proliferation (FDR 5%); accordingly, membrane, transport and intracellular organelle were enriched on the cellular component category, and neuropeptide receptor and G protein-coupled peptide receptor activities were enriched in the molecular function category. A single molecular function was enriched among the 249 genes identified in C5, zinc ion transmembrane transporter activity. Among the 1,067 genes annotated in the C8 inversion, several GO terms involved in carbohydrate metabolism, energy production and tissue remodelling, such as monocarboxylic acid transport, glucokinase activity, hexokinase activity, and glucose binding, were detected, and interestingly, several others associated with immune-related pathways, such as cytokine receptor binding and tumour necrosis factor receptor signalling (Table S6).

### Population genomics of O. edulis

#### Chromosome inversions (arrangements) distribution across the species range

Thirteen shellfish beds were analysed to explore the distribution of inversion arrangements throughout *O. edulis* range (Figure 1; Table 1). K = 2 was the most confident clustering rendered by STRUCTURE program in all shellfish beds when using the SNPs within each of the three inversions; furthermore, all individuals were easily classified as homokaryons or heterokaryons for both arrangements showing a full or intermediate composition for each cluster (Fig. 4).

**Figure 4.**
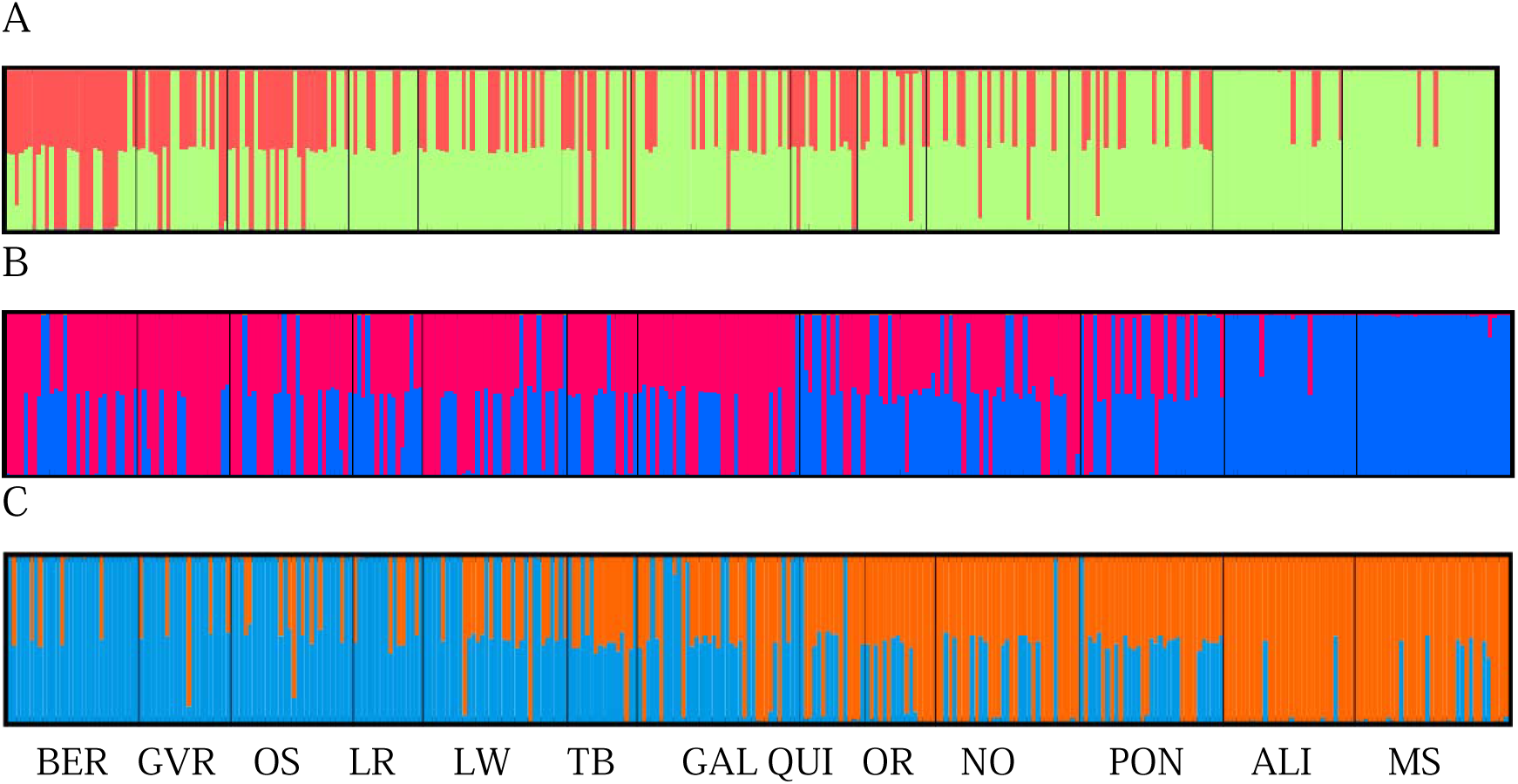
STRUCTURE analysis (K = 2) using SNPs within each of the inversions at C2 (A), C5 (B) and C8 (C) in *O*. *edulis* across the distribution range.

The distribution of arrangements for each inversion followed a consistent north-to-south cline across the species range (Fig. 5; Table S7). Both arrangements showed very similar frequencies in C5 and C8 inversions (0.500), while the “green” arrangement was the more frequent in C2 (0.732). All samples conformed to HWE at arrangment level for each of the three inversions (P > 0.05) (Table S8). Global F_ST_ across all shellfish beds was 0.294, higher for C8 (0.376) and C5 (0.311) than for C2 (0.170) (P < 0.001 in allcases). Pairwise F_ST_ showed a rough isolation by distance (IBD) north-to-south trend (P < 0.001) with notable exceptions, reflected as differential patterns between inversions, especially for C2 vs C5 and C8 (Table S9). For instance LW and LR showed no differentiation with all Atlantic Irish, French and Spanish beds for C2 (average F_ST_ = −0.006), while it was high and significant for C5 (average F_ST_ = 0.107, P < 0.001) and C8 (average F_ST_ = 0.268, P < 0.001). Accordingly, pairwise F_ST_ correlations between inversions across the distribution were low-moderate, although significant (Spearman’s rho C2 vs C5 = 0.399; C2 vs C8 = 0.358; C5 vs C8 =0.551; P < 0.001in all cases).

**Figure 5.**
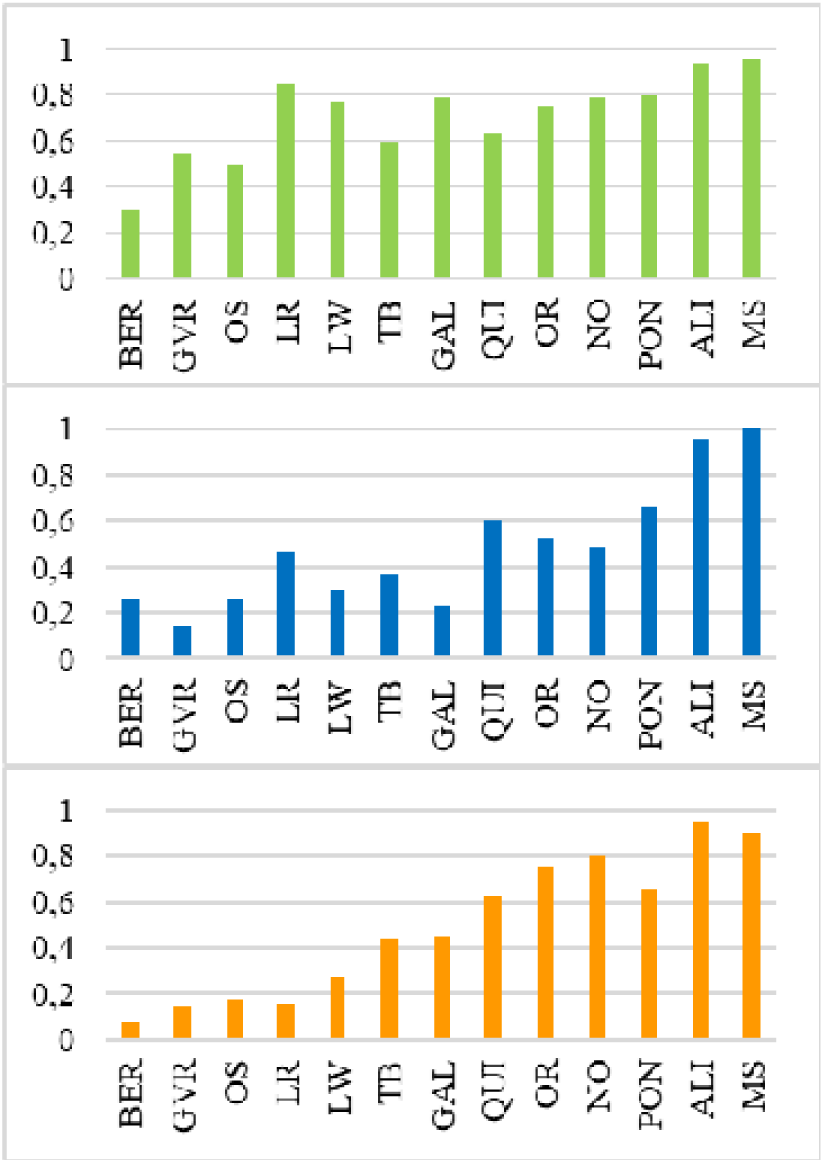
Frequency of the “south” arrangement for inversions at C2, C5 and C8; samples are ordered from north to south throughout the *O*. *edulis* distribution. Colours refer to the correspondent STRUCTURE cluster for each inversion.

Deviation from HWE was studied within each inversion suggestive of divergent selection (heterokaryon deficit) or overdominance (heterokaryon excess) using the large Irish population (GAL+LW = 384 individuals; Table 1). Significant heteroyzygote excess was only detected for C5 inversion (F_IS_ = −0.130, P = 0.012) and no deviations were detected either for C2 or C8. No significant association between pairs of inversions could be detected using exact tests for genotyping equilibrium (P > 0.05 in all the three pairwise comparisons).

#### Genetic diversity and structure of *O*. *edulis* at the light of inversions

Most beds showed high and similar genetic diversity when using the 4,741 SNPs (He ∼0.4), excluding the two Mediterranean samples, which showed lower diversity likely due to higher isolation (MS and ALI; He = 0.360 and 0.334; Table S10). Genetic diversity was slightly lower within inversions than in the rest of the genome (average He: 0.362 vs 0.398, respectively; Wilcoxon paired test, P = 0.022). Genetic differentiation (pairwise F_ST_) with the whole SNP dataset was between low and moderate (global F_ST_ = 0.055; P < 0.001; pairwise range: GRV vs OS = 0.0002 and BER vs MS = 0.1491; Table S11), the Mediterranean samples displaying higher differentiation. When excluding the SNPs within the inversions, genetic differentiation significantly decreased (F_ST_ = 0.040, P < 0.001) in accordance with the much higher differentiation detected within the inversions (total F_ST_ = 0,158; F_ST-C2_ = 0.106; F_ST-C5_ = 0,189; F_ST-C8_ = 0.166; P < 0.001 in all cases).

The most probable number of genetic clusters (K) when using the whole SNP panel was K = 3. These clusters roughly corresponded to beds from: i) the North Sea (BER, OS, GRV), ii) the Atlantic Ocean, with a gradual variation from Ireland and Scotland to France and Spain (GAL, LW, LRY, QUI, OR, NO, PO), and iii) the Mediterranean Sea (ALI, MS). Nevertheless, the influence of the chromosome inversions was remarkable, with many individuals showing an intermediate ancestry pattern throughout the distribution range (Fig. 6A). When SNPs from the three inversions were excluded, a smooth north to south variation pattern was observed across four main regions disclosed (K = 4). At both ends of the distribution range, two pure highly divergent beds were identified, Croatia (MS) and Norway (BER), and then, three differentiated groups with variable admixture degree in the south (ALI to QUI), British Isles (GAL to LR) and North Sea (OS and GRV). Higher K values were also explored (Fig. S8), and with K = 5 a connection between North Sea, TB and Cantabrian Sea was unveiled. DAPC analyses using the same SNP panels showed a similar clustering pattern across beds, especially at the ends of the distribution (Fig. S9A); however, when excluding C2, C5, and C8 inversions, beds from Spain (including QUI), British Isles and North Sea appeared further apart (Fig. S9B).

**Figure 6.**
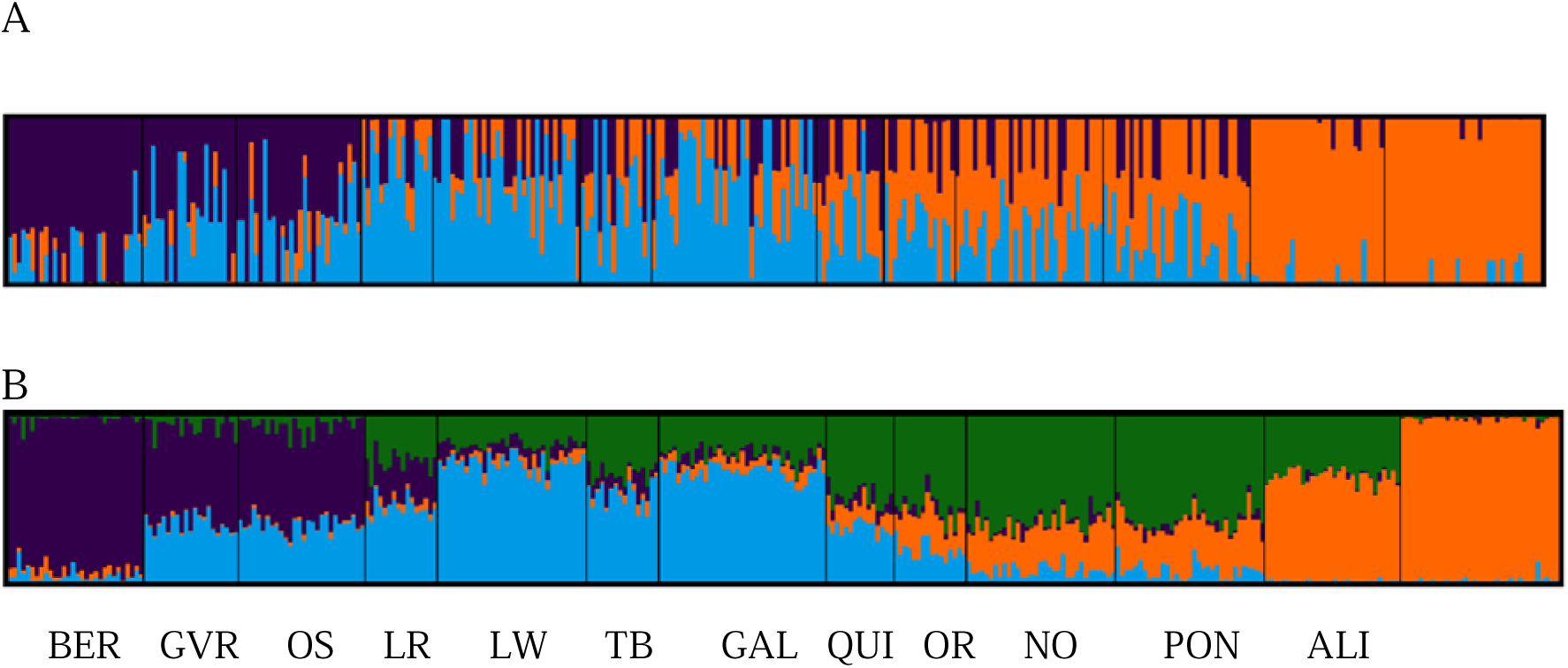
STRUCTURE analysis for *O*. *edulis* from 13 European beds using different SNP datasets: A) full SNP panel of 4,741 SNPs (K = 3); B) dataset excluding SNPs within chromosomal inversions (4,316 SNPs; K = 4).

A strong differential pattern was disclosed for the three inversions with respect to the rest of the genome when looking for outlier loci with BAYESCAN (Table 4); most SNPs within inversions were associated with divergent selection (total 76.1%; C2: 67.4%; C5: 81.0%; C8: 80.8%), while only represented 7.8% in the rest of the genome. Inversions at C5 and C8 showed a divergent selection signal much greater than C2.

**Table 4.** Outliers for divergent selection and neutral SNPs within the inversions in C2, C5 and C8 and for the rest of the genome across the 13 samples of *O*. *edulis*.

|  | C2 |  | C5 |  | C8 |  | all inversions |  | rest genome |  |
| --- | --- | --- | --- | --- | --- | --- | --- | --- | --- | --- |
|  | N | ratio | N | ratio | N | ratio | N | ratio | N | ratio |
| outliers | 97 | 1.820 | 47 | 7.231 | 168 | 4.800 | 312 | 13.851 | 338 | 0.553 |
| neutral | 47 | 0.882 | 11 | 1.692 | 40 | 1.000 | 98 | 3.574 | 4332 | 7.087 |

N: number of SNPs; ratio: number of SNPs/Mb

#### Genetic diversity and differentiation throughout the inversions

The variation in genetic diversity (He) per arrangement and genetic differentiation between arrangements (F_ST_) was inspected throughout the inversion lengths to search for insights into the evolutionary forces operating on these genomic regions. The analysis was made in the 13 samples throughout the flat oyster distribution and in the big Irish sample using single SNPs but also using sliding windows averaging He and F_ST_ estimators (Figure S10; Tables S12 and S13). When using the 13 samples, much lower genetic diversity was observed within each arrangement for the three inversions than in the rest of the genome (C2: He_red_ = 0.065 and He_green_ = 0.160; C5: He_pink_ = 0.108 and He_darkblue_ = 0.103; C8: He_blue_ = 0.159 and He_orange_ = 0.204; He_WG_ ∼0.400). Furthermore, the F_ST_ between arrangements was much higher (F_STC2_ = 0.681; F_STC5_ = 0.609; F_STC8_ = 0.494; P < 0.001 in all cases) than the global F_ST_ in the whole genome excluding inversions (F_STWG_ = 0. 040). These figures were also reflected by the high number of SNPs fixed or nearly fixed for alternative variants in both arrangements (F_ST_ > 0.9: C2 = 34.4%; C5 = 34.8%; C8 = 16.2%).

All data support a strong divergent selection operating on these genomic regions. Interestingly, the pattern was rather different in the three cases, not only for the lower differentiation between arrangements in C8 inversion but also for the specific F_ST_ profiles across the inversion length. C5, for instance, showed a strong differentiation close to the 5’ end breakpoint and to a minor extent to the 3’ end following a rough valley-profile; conversely, both C2 and C8 displayed saw-peak profiles around multiple highly differentiated subregions. As expected, these F_ST_ profiles are nearly identical to the LD profile (Figure S4) and suggest selective sweeps around specific subregions in the three inversions. When using the big Irish sample results were very similar (Fig. S10; Table S12).

The genes close to the breakpoints of inversions (< 100 kb) and around the putative adaptive peaks within each inversion were inspected to identify potential candidates or functions explaining the high differentiation observed (Table S14). It is worth noting the important number of low quality or uncharacterized proteins in the current annotation (∼40%), which hampered the analysis and underscores the need for improving annotation in mollusc genomes. Two notable genes were detected at the 3’ end breakpoints of C5 and C8: putative inhibitor of apoptosis and Sushi nidogen and EGF-like domain-containing protein 1, involved respectively in cell death and extracellular matrix constitution. Both apoptosis and extracellular matrix have been identified as key functions showing differentially expressed genes in flat oyster in response to *B. ostreae* (Ronza et al., 2018). Important genes related to energy production, digestion, growth, reproduction and immunity, such as cyanocobalamin reductase, E3 ubiquitin-protein ligase TRIM33-like, Ig-like and fibronectin type-III domain-containing protein 2, membrane progestin receptor beta-like, mitogen-activated protein kinase 1-like, pancreatic lipase-related protein 2-like, somatostatin receptor type 2-like, T-cell acute lymphocytic leukemia protein 1-like, and toll-like receptor 4 were detected around the adaptive peaks inspected. One specific cluster in C8 (62,793,855 bp) included several genes involved in spermatogenesis and growth (spermatogenesis-associated protein 24-like, growth hormone-regulated TBC protein 1-A-like, mitotic checkpoint protein BUB3-like and sex peptide receptor-like).

## Discussion

The genetic structure of flat oyster has been intensively studied since the 1990s using allozyme, microsatellite, and SNP markers, which reflects the importance of this species from ecological and production perspectives (Zu Ermgassen et al., 2023). These studies consistently reported four main genetic groups geographically distributed throughout its native range, namely Mediterranean, south Atlantic, British Isles and North Sea (Vera et al., 2016; Pouvreaeu et al., 2023). It has been suggested that human-mediated movements related to production have blurred some areas of its native distribution (Lapègue et al., 2022; Monteiro et al., 2024). However, the most recent studies using SNP genomic screening have provided new relevant insights associated with phylogeographic events and adaptation to biotic or abiotic environmental factors (Vera et al., 2019; Sambade et al., 2022; Kamermans et al., 2023; Monteiro et al., 2024; Robert et al., 2025). Of note within this landscape, the identification of big chromosome inversions in three chromosomes related to historical events that could play specific adaptive roles (Sambade et al., 2022, Monteiro et al., 2024).

Using a low-density SNP array, selected from the previous flat oyster ∼14K array reported by Gutierrez et al. (2017), we reanalysed the genetic structure of flat oyster, focusing on the origin and adaptive role of inversions. The SNP array developed and applied in our study enabled robust genotyping (missing data ∼1.5%) across the whole genome (∼5 SNPs / Mb). The MAF > 0.05 threshold used for filtering SNPs from the 14K array likely explains the higher genetic diversity in our study (He ∼ 0.390) compared to previous studies (He ∼ 0.310; Vera et al., 2019; Sambade et al., 2022).

While inversions have been found across many taxa, mapping and characterizing inversion breakpoints remains challenging, often due to the presence of complex tandem repeats and transposable elements (TEs) (Puerma et al., 2014; Villeoutreix et al., 2021). The use of the 5K array in a big flat oyster sample (384 individuals) enabled us a confident LD estimation across the whole genome, confirming the three inversions previously reported. Furthermore, the breakpoints of the C8 inversion, and at the 5’ and 3’ ends of C2 and C5 inversions, respectively, could be precisely mapped and validated through PCR assays. The failing at the 5’ end at C5 and at the 3’ end of C2 breakpoints could be related to the positional effect on crossing over in the adjacent chromosomal regions, as reported in other species (Korunes and Noor, 2019; Koury, 2023). The PCR here used for validating inversion breakpoints might be used as a molecular tool for inversion genotyping, but it cannot distinguish between homokaryon and heterokaryon genotypes, thus representing a dominant marker. However, the large number of loci fixed for alternative alleles in the three inversions could be easily applied to design a multiplex SNaPshot or MassARRAY covering the entire inversion length for genotyping to address relevant features, such as double recombinant events (Berdan et al., 2023).

The three inversions appear to be paracentric according to flat oyster karyotype (Thiriot-Quiévreux, 1984) and LD information, and span a substantial portion of the flat oyster genome (∼8%). Long paracentric inversions are common in other species (Kirkpatrick, 2010; Wellenreuther and Bernatchez, 2018). Additionally, both arrangements were found at high frequency when considering the whole flat oyster distribution (0.5/0.5 for C5 and C8; 0.25/0.75 for C2), indicating high heterokaryon frequencies in many samples. Paracentric inversions give rise to very abnormal acentric and dicentric chromosomes during meiosis I if single crossing-over occurs, representing a strong mechanism for disrupting recombination. This would reduce heterokaryon fertility, unless some mechanism removing abnormal gametes, such as meiotic drive or gamete segregation to polar bodies are operating (Sturtevant and Beadle, 1936; Koury, 2023; Samura et al., 2023; Zakharov, 2024). We did not find significant deviations from HW proportions in the 13 flat oyster samples across the distribution range, but instead a significant heterokaryon excess (P = 0.022) in C5 inversion in the big Irish sample and a slightly low global negative F_IS_ in the 13 natural beds studied. Overdominance has been claimed and demonstrated in some cases as a mechanism for maintaining inversions (Faria et al., 2019; Knief et al., 2017). Regardless of the mechanisms underlying the maintenance of these polymorphisms in flat oyster, our data suggest no negative fitness effects on heterokaryons for the three inversions.

Identifying markers, genes, and functions associated with inversions is an essential starting point to deepen into their origin, maintenance and distribution throughout the flat oyster natural range. The persistence of inversions across multiple species and populations suggests they may confer adaptive advantages. From an evolutionary perspective, the conservation of certain inversions over time indicates that they may be subject to selection providing beneficial allelic combinations for adaptation to specific environments (Faria et al., 2019; Füller et al., 2020). The C2 and C8 inversion lengths were notably longer in our study than that reported by Monteiro et al. (2024), likely due to the different genome assemblies used as reference in both studies, Gundappa et al. (2022) and Boutet et al. (2022), respectively. The established breakpoints and the LD variation observed across the whole length of the three inversions in our study fits well to the assemblage reported by Gundappa et al. (2022) and to the chromosome size of C2, C5 and C8 in the new chromosome-level assembly reported by Li et al. (2023). Therefore, discrepancies might be related to assembly issues or the presence of important structural variants in the different individuals used for assembly.

We identified specific repetitive elements (REs) flanking the four mapped breakpoints at C2, C5 and C8 inversions (overlapping in C5, and between 1.3 and 9.4 kb for C8 and C2): several LINE type elements on one side in all cases and DNA/hAT or PLE/Naiad elements on the other. Penelope-like, LINE and DNA/hAT elements have been suggested in the origin of inversions in *Drosophila* and other species (Evgen’ev et al., 2000; Lee et al., 2008; Araya et al., 2025). These three elements represent an important RE proportion in the three chromosomes (∼14% each element), where REs constitute around 30% of the total chromosome length, so we cannot discard that our observation could occur just by chance. We also identified short tandem repeated sequences at both ends of C2 and C5, which could be related to staggering associated with a break and joining mechanism (Villoutreix et al., 2021).

The genetic structure of flat oyster was explored at the light of the three inversions, both to enrich previous observations for managing this valuable resource, as well as for understanding the role that inversions may play in flat oyster evolution and adaptation. The main genomic regions identified across the distribution range in our study are in accordance with those recently reported by Monteiro et al. (2024), so a consistent and valuable information for sustainable management of wild and production areas is available from these studies. Regarding the population genomic features of inversions, Monteiro et al. (2024) suggested a phylogeographic explanation involving two divergent lineages in the north and the south of flat oyster distribution followed by a secondary contact, without precluding some current adaptive role. Indeed, the frequency distribution of arrangements observed in both studies for the three inversions follows a significant north to south cline, either including or excluding the Mediterranean samples, which might fit to the phylogeographic hypothesis. However, several insights suggest a role in adaptation when comparing their distribution pattern and population parameters with respect to the rest of the genome. Firstly, despite the clinal variation observed, the correlation between pairwise F_ST_ across the 13 samples between inversions was significant, but low (especially C2 vs C5/C8 ∼0.32) or moderate (C5 vs C8 ∼0.55). Secondly, SNP markers within inversions showed much higher pairwise F_ST_ differentiation than the rest of the genome and in fact the STRUCTURE plot obtained when including and excluding inversions was quite different. As previously suggested, inversions and low-recombining regions differentiate populations more strongly than collinear recombinant regions, either along an ecogeographic cline or at a fine-grained scale (Mérot et al., 2020). Thirdly, most outliers detected across the distribution range of flat oyster were located within inversions strongly supporting divergent selection associated to those genomic regions. This pattern of selection is likely not recent (see below) and could underlie the existence of two old lineages. However, these polymorphisms might still be under selection in the current conditions or be harnessed as an important source of variation to adapt to the specific biotic and abiotic conditions of the current flat oyster distribution.

As outlined above, several observations suggest an ancient origin of these inversions in flat oyster. Inversions represent unique mutational events that are usually removed from populations due to lower fertility of heterozygotes, unless a significant fitness benefit, either from positive selection or overdominance, might counterbalance this fact (Berdan et al., 2023). The new arrangement lacks genetic diversity and accordingly the older arrangement has been usually associated with higher genetic diversity for recent events (Kapun et al., 2023). Genetic diversity in the new arrangement increases through mutation, gene-flux (double crossing-over) and gene conversion (Charlesworth, 2023). In our study, both arrangements at the three inversions showed substantial genetic diversity suggesting quite old events; however, the “orange” arrangement of C8 (He = 0.204) and the “green” one of C2 (He = 0.160) showed significantly higher genetic diversity than their “blue” (He = 0.159) and “red” (He = 0.065) counterparts, respectively, suggesting the southern arrangements to be older. The ancient origin of flat oyster inversions is also strongly supported by the high genetic differentiation between both arrangements (F_ST_ ≥ 0.50) with respect to the global differentiation across samples when excluding markers within inversions (F_ST_ < 0.04). This aligns well with the high number of outliers and extreme F_ST_ detected for SNPs within inversions supporting strong divergent selection in their origin and evolution.

The analysis of F_ST_ and LD across the inversions unveiled specific patterns of differentiation for the three inversions suggesting selective sweeps around several focal SNPs. Coalescence models on the evolution of inversions suggest that selection increases divergence by hitchhiking neutral loci linked to target sites (Charlesworth, 2023). We found heterogeneous patterns of F_ST_ for the three inversions, with peaks distributed across the whole inversion length in C2 and C8, but closer to both ends in C5. The latter pattern could be related to a positional effect of the breakpoint on gene expression of adjacent genes (Kirkpatrick, 2010). We could identify some genes or clusters potentially associated with adaptation around those peaks, some of them related to reproduction such as membrane progestin receptor beta-like, spermatogenesis-associated protein 24-like, and sex peptide receptor-like. Reproduction related genes including in inversions have been associated to divergent selection and even speciation (Berdan et al., 2023).

Functional enrichment of the genes annotated within inversions unveiled specific functions that could be associated with adaptation to a diverse environment, including abiotic (temperature and salinity) and biotic (pathogens, intra and interspecific competition) factors. The C2 inversion encompassed notable functional diversity, including neuropeptide signalling pathways, epigenetic regulation, immune response, and metabolic processes. This suggests a putative role of this inversion for resilience to environmental challenges. Genes related to neuropeptide signalling and external stimulus detection along those related to transmembrane transport and ion channel activity suggest potential adaptation to temperature and salinity variation (Clark et al., 2013, 2020; Signorini et al., 2025). Also, genes associated with ciliary activity and phytoplankton filtration points toward differences in feeding strategies and energy metabolism (Marinković et al., 2019). In contrast, the inversion on chromosome 5 included genes primarily associated with zinc transport and homeostasis; Zn represents an essential ion for numerous biological functions, including enzymatic activity, gene expression regulation, and oxidative stress response. Differential expression of genes associated with this function has been involved in response to Zn pollution in *Magallana gigas* (Meng et al., 2021) and *Crassostrea hongkongensis* (Li et al., 2020).

The C8 inversion has been previously reported to be associated with resistance to *B*. *ostreae*. The region delimited in our study includes 14 of the 16 resistance-associated outliers previously reported (Vera et al., 2019; Sambade et al., 2022; Kamermans et al., 2023). The genes within this inversion were enriched in functions such as carbohydrate metabolism, energy production, tissue remodelling, and immune pathways, including cytokine receptor signalling and tumour necrosis factor signalling (Canesi et al., 2022). This suggests that the region may influence immune response and tissue repair processes following infections by *B*. *ostreae*, an intracellular parasite. Notably, naïve or low affected samples such as Bergen and Lough Swilly, displayed a higher frequency of the “blue” arrangement than their geographically close affected counterparts, Oosterschelde and Galway, respectively. The high frequency of the “orange” arrangement in the south and the association of *Bonamia* with warmer temperatures (Audemard et al., 2008) could explain the association observed. However, a spurious correlation due to the north to south clines of arrangements and the latitudinal distribution of the parasite could also explain the association observed at population level. Further analyses should be carried out to confirm the association observed at individual level for its putative application on controlling bonamiosis.

## Conclusions

We precisely mapped the three big chromosome inversions of flat oyster as an essential starting point to delve into their population and evolutionary meaning. The three inversions span a significant portion of the flat oyster genome and seem to represent old mutational events. Although a north-south cline was unveiled for the three inversions, several insights suggest specific adaptive roles in the current flat oyster distribution. The strong divergent selection between arrangements followed specific patterns for each inversion associated to subsets of genes involved in putative roles related to feeding, stress resilience, energetic resources and resistance to pathogens. The information obtained will aid for more deepen studies aimed at understanding their adaptive role to be applied for sustainable conservation of this emblematic resource.

## Acknowledgements

We acknowledge the Centro de Supercomputación de Galicia (CESGA) for providing computational resources and support, which were essential for the analyses conducted in this study.

We also thank Dr. Daria Ezeta Balić, Dr. Ivana Buselic and her colleagues for kindly providing oyster samples that were part of their work on the Adriatic coast.

## Notes

### Competing Interest Statement

The authors have declared no competing interest.

